# Universal protection of allogeneic cell therapies from natural killer cells via CD300a agonism

**DOI:** 10.1101/2024.05.05.592600

**Authors:** Shu-Qi Zhang, Faith Thomas, Justin Fang, Kathryn Austgen, Chad Cowan, G. Grant Welstead

## Abstract

Immunogenicity limits the persistence of off-the-shelf, allogeneic cell therapies and transplants. While ablation of human leukocyte antigen (HLA) removes most T cell and humoral alloreactivity, no solution has enabled universal protection against the resulting natural killer (NK) cell response. Here, we engineered Trans Antigen Signaling Receptors (TASR) as a new class of NK inhibitory ligands and discovered CD300a, a previously inaccessible receptor, as a functional target. CD300a TASR outperformed leading alternative strategies in focused screens, including CD47 and HLA-E, and was solely capable of universally protecting allogeneic T cells against a large human cohort (45/45 donors), spanning diverse demographics and NK cell phenotypes. A model allogeneic T cell therapy co-expressing an anti-CD19 Chimeric Antigen Receptor (CAR) and CD300a TASR, produced using multiplexed non-viral integration, exhibited enhanced B cell killing potency under allogeneic immune pressure. CD300 TASR represents a universal solution to NK alloreactivity, broadening the population that could be effectively treated by next-generation allogeneic cell therapies.

**Category:** Immunobiology and Immunotherapy

**Key Points:**

- An engineered CD300a agonist ligand (CD300a TASR) universally protects HLA-deficient allogeneic T cells from NK-mediated rejection.
- CD300a TASR is more protective in CMV seropositive hosts than HLA-E ligand and enhances CAR-T efficacy under allogeneic immune pressure.

## Introduction

Off-the-shelf cellular therapies, such as T and CAR-T cells, promise to build on the success of its autologous counterparts while providing patients a well-defined, scalable, and cost-efficient drug product that can be administered on-demand^1^. Host immune rejection, however, is a major obstacle that limits their persistence, efficacy, and redosing potential^2^. While elimination of donor HLA class I (HLA-I) is a common strategy to evade T cell alloreactivity and HLA-specific anti-donor antibodies (ADAs), this strategy also unleashes NK alloreactivity owing to loss of inhibitory signaling from Killer cell immunoglobulin-like receptors (KIR) binding to HLA-I that normally restrains the NK cell response^3,4^. Current approaches to reinstate NK inhibition rely on the expression of natural cloaking ligands such as HLA-E^3,5,6^, HLA-A^7^, HLA-C^8^, and CD47^9^ to agonize NKG2A, KIR, and SIRPα inhibitory receptors. These targets, however, are only expressed on limited subsets of NK cells or upon sustained cytokine stimulation, rendering the ligands ineffective against hosts with low frequencies of the corresponding NK subset^3^. In this study, we discover and assess a universal solution against NK alloreactivity to enhance the persistence and efficacy allogeneic T cell therapies against all hosts.

## Methods

The objective of this study was to assess the function of CD300a TASR and alternative ligands for protecting allogeneic T/CAR-T cells from rejection by human primary NK cells and peripheral blood mononuclear cells (PBMCs), and to identify demographic and phenotypic features that affect ligand function. T and CAR-T cells are challenged with NK and PBMCs for 20 or 72 hours, respectively, and then subjected to small molecule or antibody-based fluorescent barcoding flow cytometry described previously^10,11^. For competition assays, pooled T cell targets are co-cultured with alloreactive T (allo-T) and/or NK cells for 20 hours prior to conventional flow cytometry. See supplemental methods for detailed cell isolation and assay protocols.

Human primary T cell targets are engineered for knock-out (KO) by CRISPR/Cas9 and knock-in (KI) by non-viral homology-directed repair (HDR) in accordance with previous protocols^12^. HLA-A2 allo-T cell clones are generated by orthotopic TCRαβ replacement as previously described^13^. For mRNA-based screening, 1-2 million activated T cells with Beta-2-Microglobulin (B2M) KO were electroporated (EP) in 20 µL P3 buffer containing 1.5 ug of mRNA using code CM-137 and rested overnight. Transgene expression was verified by staining with relevant antibody and/or recombinant protein ligand and acquired on flow cytometry. See supplemental methods for detailed T cell engineering, culture, and staining protocols.

## Results and Discussion

We first developed an *ex-vivo* model of cellular rejection by measuring the survival of human allogeneic T cell targets co-cultured with human primary cell effectors using a modified fluorescently barcoded flow cytometry readout (**Supplemental Fig. 1a**)^10^. This method enabled sensitive and high-throughput measurement by direct counting across multiple E:T ratios with high reproducibility in IC50 value (**Supplemental Fig. 1b**,**c**). Ablation of HLA-I expression on T cells via functional KO of B2M prevented rejection by allo-T cells, but consistently introduced NK cell alloreactivity across multiple donor replicates, confirming missing-self mechanism of NK recognition in this context (**Supplemental Fig. 2**)^4^.

We applied this experimental system with mRNA-mediated expression and CRISPR-mediated KO to rapidly screen prior “cloaking” strategies to inhibit NK cells and identified HLA-E as the sole potent ligand (**Fig. 1a, Supplemental Fig. 3a**)^3,5,6^. We hypothesized that surface expression of single-chain antigen binding fragment (scFV) on an inert scaffold, termed TASRs, can agonize a wider repertoire of NK inhibitory receptors than is currently possible. Using a CD8 hinge and transmembrane as the initial scaffold, we identified a NKG2A and a CD300a binding TASR that functioned in a dose-dependent manner (**Fig. 1a, Supplemental Fig. 3a-c**). NKG2A TASR agonizes the same target as HLA-E, while CD300a is an unexplored target for this application. Unlike NKG2A, CD300a is a classical inhibitory receptor known to be expressed on all NK cells, and signals through immunoreceptor tyrosine-based inhibitory motifs (ITIMs) upon binding phosphatidylserine (PS) and phosphatidylethanolamine (PE) on the surface of apoptotic cells (**Fig. 1b**)^14,15^.

**Figure 1:**
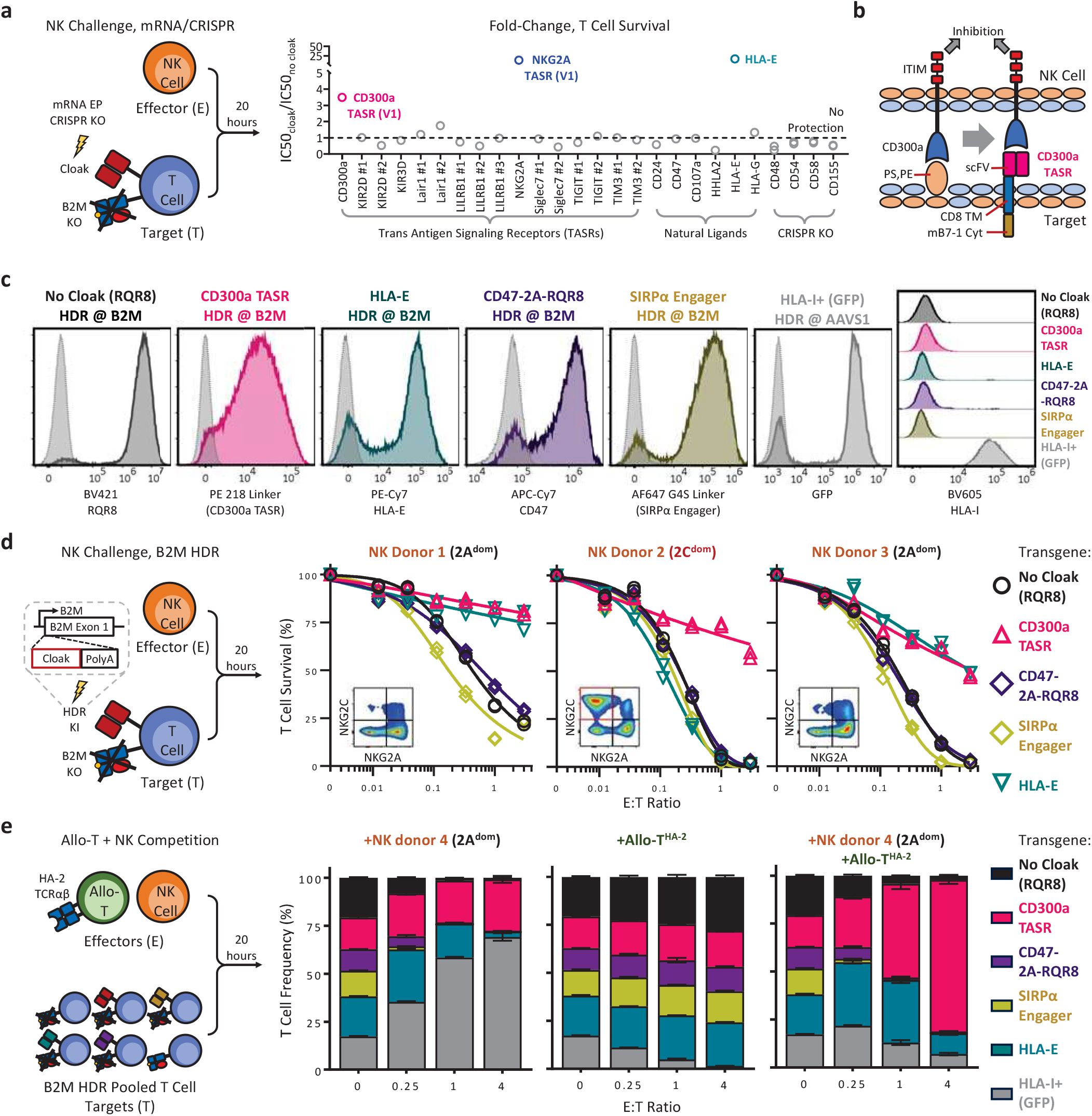
Discovery and Assessment of CD300a TASR. a, NK challenge assay to screen cloaking strategies on B2M KO T cells. Ligands are expressed by mRNA EP, and KO by CRISPR/Cas9. Y-axis shows ratio of IC50 value of the indicated cloaked T cell over uncloaked negative control. IC50 represents E:T ratio corresponding to 50% target T cell survival, derived from one 6–7-point curve of T cell survival at various E:T ratios challenged with one NK cell donor, with higher values indicating enhanced persistence. N = 1 technical replicate per condition. b, CD300a TASR design and mechanism of action. c, phenotype by flow cytometry of engineered T cells expressing the indicated transgenes at the target locus by non-viral HDR, gated single cell lymphocytes. grey dotted histograms indicate negative control T cells stained with the same markers. CD47 is endogenously expressed on T cells, and so the RQR8 epitope tag was co-expressed using 2A self-cleaving peptide to facilitate purification transgene-expressing cells. d, NK challenge assay with B2M KO T cells expressing cloaking transgene from the B2M locus from (c), challenged with three NK cell donors. Inset shows the NK cell phenotype by flow cytometry, gated CD3-CD56+. N= 2 technical replicate curves per condition. e, Competition assay of pooled T cell targets from (c) co-cultured with an allo-T cell clone and/or NK cell effector. Allo-T cells express TCRαβ specific for minor histocompatibility antigen 2 (HA-2) peptide presented on HLA-A2. Target T cells are serotyped HLA-A2^+^, and effector cells are HLA-A2^-^. Y-axis shows frequency of the indicated T cell member after challenge with indicated effector cell. N = 3 technical replicates per condition.

We set out to optimize the expression and function of CD300a TASR version 1 by testing several mutant constructs using our mRNA screening platform. We discovered a mouse B7-1 transmembrane (TM) and cytoplasmic (Cyt) domain that enhanced expression and an inverse correlation between TASR hinge length and the function of CD300a TASR but not NKG2A TASR (**Supplemental Fig. 4, 5a-e**)^16^. An optimized version 2, containing no hinge but stabilized by the mouse B7-1 cytoplasmic tail, and hereafter termed CD300a TASR, exhibited significantly enhanced function while maintaining a fully human extracellular domain (**Supplemental Fig. 5f**,**g**).

CD300a TASR outperformed the current best-in-class ligands HLA-E and CD47, as well as the recently published SIRPα and TIM3 engagers in NK protection^3,6,9,17^. To provide head-to-head comparisons, we inserted each ligand into the B2M locus of human T cells concomitant with B2M KO by non-viral HDR under constitutive expression by the endogenous B2M promoter and confirmed surface expression (**Fig. 1c**). Challenged with NK cells, CD300a TASR and HLA-E outperformed CD47 and SIRPα engager, while CD300a TASR was the only protective ligand against a donor with high frequencies of an NKG2A^-^NKG2C^+^ NK cell phenotype, hereby termed 2C^dom^ (**Fig. 1d**)^3^. The dominance of CD300a TASR over CD47 was observed under all conditions tested, including when overexpressed via mRNA, inserted into the AAVS1 locus via HDR under EF1α promoter, and challenged with NK cells cultured in high-dose IL-2, which upregulates SIRPα (**Supplemental Fig. 6**)^17^. Using mRNA EP, we further determined that CD300a TASR outcompeted TIM3 engager against 4/4 NK donors under comparable levels of expression (**Supplemental Fig. 7**).

As an orthogonal readout, we pooled HDR-edited T cells from the previous experiment along an HLA-I+ control and challenged with allo-T and NK cells in a competition assay. The identity and frequency of each T cell member was measured by conventional multicolor flow cytometry (**Supplemental Fig. 8a-e**). As expected, CD300a TASR emerged as the dominant survivor when challenged with both NK and allo-T cells (**Fig. 1e**). This result was reproducible with a second 2C^dom^ NK cell donor and alloreactive T cell clone of different specificity, as well as any combination thereof (**Supplemental Fig. 8f**). Thus, CD300a TASR with HLA-I ablation is solely capable of protecting against both T and NK cell alloreactivity across all donors.

We next assessed CD300a TASR and HLA-E, the most potent alternative, against a large human cohort to assess the heterogeneity of responses across donors and to identify correlates of protection. To model the physiological context of intravenously injected T cell therapies, we measured the survival of cloaked T cells challenged with PBMCs (**Fig. 2a, Supplemental Fig. 9**). CD300a TASR and HLA-E were expressed at comparable levels from the AAVS1 locus based on RQR8 expression (**Fig. 2b**)^18^. 45 PBMC donors were selected for diversity in ethnicity, age, gender, and CMV seropositivity due to their known effects on NK cell function and phenotype (**Fig. 2a**) ^4,19–21^. While large variations in CD57, KIR, NKG2A, and NKG2C NK marker expression was observed, CD300a was uniformly expressed on NK cells in all donors (**Supplementary Fig. 10**). Within this cohort, CD300a TASR protected B2M KO T cells against all PBMC donors tested (45/45), indicating universal protection against NK cell reactivity (**Fig. 2c-d, Supplemental Fig. 11**). CMV serostatus emerged as the most significant demographic determinant of reduced protection by HLA-E and correlated with the frequency of the well-known adaptive NK cell subset (**Fig. 2e, Supplemental Fig. 10a**,**d**)^20^. While HLA-E lost protection with increasing frequencies of adaptive NK cells, CD300a TASR maintained protection (**Fig. 2f**). Since more than half the human population are CMV seropositive, we expect the use of CD300a TASR in allogeneic cell therapies to significantly expand the addressable patient population^22^.

**Figure 2:**
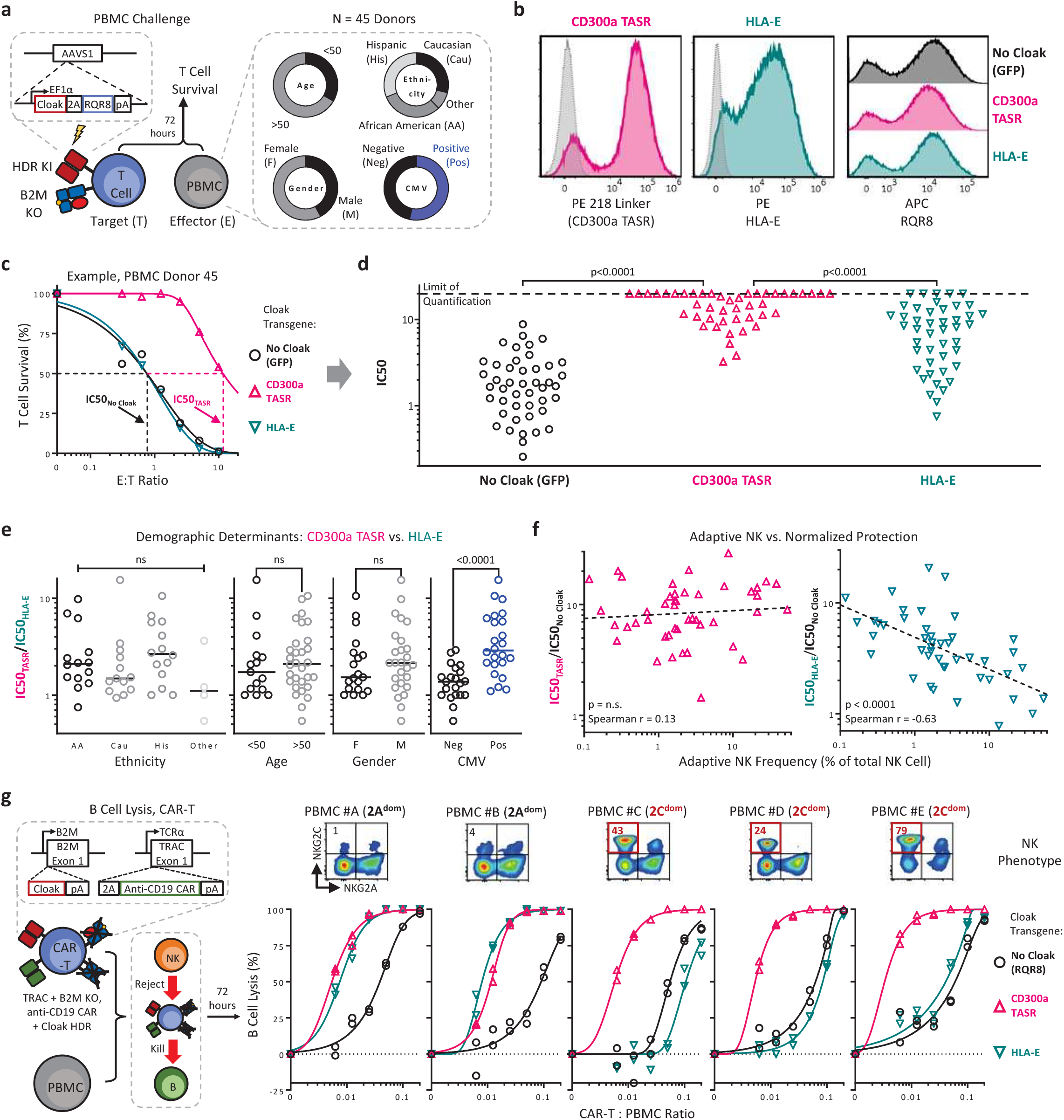
CD300a TASR universally protects against NK cells and enhances CAR-T functional potency. a, Study design and demographic overview of the PBMC donors used. T cells are engineered to co-express cloaking transgene and RQR8 via 2A self-cleaving peptide under control of EF1α promoter by non-viral HDR. b, Phenotype by flow cytometry of the three engineered T cell targets, gated on live single lymphocytes. Label indicates cloaking transgene. grey dotted histograms indicate negative control T cells stained with the same markers. c, Example of PBMC challenge assay result and derivation of IC50 values. d, Aggregate results as in (c) against 45 PBMC donors. Each datapoint represents IC50 value of the indicated cloaking ligand against PBMCs from one donor. Limit of quantitation in IC50 set to be twice the highest E:T ratio used. Wilcoxon matched pairs signed rank test. e, Association of PBMC donor demographics from (a) with functional data from (d), Kruskal-Wallis for Ethnicity, Mann-Whitney for other. Y-axis represents ratio of IC50 between CD300a TASR and HLA-E cloaking ligand, with one indicating equal protection. f, Relationship between the adaptive NK cell frequency of the PBMC donor (phenotype NKG2A^-^NKG2C^+^CD57^+^CD56^lo^CD16^hi^) and functional potency of CD300a TASR (left) and HLA-E (right), normalized to non-cloaked control from (d). Dotted line represents linear fit of log-log transformed data. N = 45 PBMC donors. g, (bottom row) B cell lysis assay for engineered CAR-T cell therapy containing the indicated cloaking transgene against the indicated PBMC donors. N = 2 technical replicate curves per condition. (top row) phenotype of NK cells from the respective PBMC donor.

Lastly, we integrated recent innovations to produce a model allogeneic CAR-T cell therapy co-expressing a clinically relevant anti-CD19 CAR into the TRAC locus and cloaking ligand into the B2M locus via multiplex non-viral HDR (**Supplemental Fig. 12a**,**b**)^12,23^. Expression of CD300a TASR, HLA-E, and RQR8 control cloaking ligand in CAR-T cells was confirmed (**Supplemental Fig. 12c**). To model CAR-T function under a physiological allogeneic environment, we dosed our engineered CAR-T cells into PBMCs and measured killing of CD19-expressing B cells under NK alloreactivity (**Fig. 2g, Supplemental Fig. 12d-h**). CD300a TASR enhanced B cell killing potency in all PBMC donors (5/5) and outcompeted HLA-E in all donors with 2C^dom^ NK phenotype (**Fig. 2g**). This result was not due to intrinsic differences, as all CAR-T cells had similar potency against CD19-expressing Raji cells in the absence of allogeneic pressure (**Supplemental Fig. 13a**). In addition, TASR-mediated protection against NK cells was maintained in the presence of CAR (**Supplemental Fig. 13b**). We expect the B cell lysis model to be relevant for allogeneic CAR-T therapy against systemic lupus erythematosus (SLE), where B cell depletion is the main mechanism of action for dampening the autoreactive antibody response^24^.

In summary, we have demonstrated that CD300a TASR acts as a universal ligand against NK cell alloreactivity. Its combination with HLA and TCRαβ ablation to prevent T cell alloreactivity, ADAs, and GvHD paves the way for fully immuno-evasive CAR-T cells that are effective for anyone in need of treatment, from cancer to autoimmunity and beyond.

## Acknowledgements

We thank Chantal Kuhn for stimulating discussions and for reviewing the manuscript, and Zhenyu Luo for providing the luciferase-expressing Raji cell line.

## Authorship Contributions

S-Q.Z. conceived and developed the TASR modality. S-Q.Z. and C.C. conceived and designed the study. S-Q.Z, F.T., and J.F. designed, performed, and analyzed data for all experiments. S-Q.Z., K.A., C.C., and G.G.W. supervised the study. S-Q.Z. and G.G.W. wrote the manuscript with feedback from all authors.

## Conflict of Interest Disclosure

All authors are current or former employees of Clade Therapeutics. C.C is a founder of Clade Therapeutics. A patent application has been filed by Clade Therapeutics for the technology described here.

## References

1. Depil S, Duchateau P, Grupp SA, Mufti G, Poirot L. ‘Off-the-shelf’ allogeneic CAR T cells: development and challenges. Nat. Rev. Drug Discov. 2020;19(3):185–199.

2. Turtle CJ, Hanafi L-A, Berger C, et al. CD19 CAR–T cells of defined CD4+:CD8+ composition in adult B cell ALL patients. J Clin Invest. 2016;126(6):2123–2138.

3. Jo S, Das S, Williams A, et al. Endowing universal CAR T-cell with immune-evasive properties using TALEN-gene editing. Nat. Commun. 2022;13(1):3453.

4. Anfossi N, André P, Guia S, et al. Human NK Cell Education by Inhibitory Receptors for MHC Class I. Immunity. 2006;25(2):331–342.

5. Gornalusse GG, Hirata RK, Funk S, et al. HLA-E-expressing pluripotent stem cells escape allogeneic responses and lysis by NK cells. Nat Biotechnol. 2017;35(8):765–772.

6. Degagné É, Donohoue PD, Roy S, et al. High-specificity CRISPR-mediated genome engineering in anti-BCMA allogeneic CAR T cells suppresses allograft rejection in preclinical models. Cancer Immunol. Res. 2024;

7. Furukawa Y, Ishii M, Ando J, et al. iPSC-derived hypoimmunogenic tissue resident memory T cells mediate robust anti-tumor activity against cervical cancer. Cell Rep. Med. 2023;4(12):101327.

8. Xu H, Wang B, Ono M, et al. Targeted Disruption of HLA Genes via CRISPR-Cas9 Generates iPSCs with Enhanced Immune Compatibility. Cell Stem Cell. 2019;24(4):566-578.e7.

9. Hu X, Manner K, DeJesus R, et al. Hypoimmune anti-CD19 chimeric antigen receptor T cells provide lasting tumor control in fully immunocompetent allogeneic humanized mice. Nat. Commun. 2023;14(1):2020.

10. Krutzik PO, Clutter MR, Trejo A, Nolan GP. Fluorescent Cell Barcoding for Multiplex Flow Cytometry. Curr. Protoc. Cytom. 2011;55(1):6.31.1-6.31.15.

11. Junker F, Teixeira PC. Barcoding of live peripheral blood mononuclear cells to assess immune cell phenotypes using full spectrum flow cytometry. Cytom. Part A. 2022;101(11):909–921.

12. Shy BR, Vykunta VS, Ha A, et al. High-yield genome engineering in primary cells using a hybrid ssDNA repair template and small-molecule cocktails. Nat Biotechnol. 2022;1–11.

13. Schober K, Müller TR, Gökmen F, et al. Orthotopic replacement of T-cell receptor α-and β-chains with preservation of near-physiological T-cell function. Nat. Biomed. Eng. 2019;3(12):974–984.

14. Zenarruzabeitia O, Vitallé J, Eguizabal C, Simhadri VR, Borrego F. The Biology and Disease Relevance of CD300a, an Inhibitory Receptor for Phosphatidylserine and Phosphatidylethanolamine. J Immunol. 2015;194(11):5053–5060.

15. Cantoni C, Bottino C, Augugliaro R, et al. Molecular and functional characterization of IRp60, a member of the immunoglobulin superfamily that functions as an inhibitory receptor in human NK cells. Eur. J. Immunol. 1999;29(10):3148–3159.

16. Lin Y-C, Chen B-M, Lu W-C, et al. The B7-1 Cytoplasmic Tail Enhances Intracellular Transport and Mammalian Cell Surface Display of Chimeric Proteins in the Absence of a Linear ER Export Motif. PLoS ONE. 2013;8(9):e75084.

17. Gravina A, Tediashvili G, Zheng Y, et al. Synthetic immune checkpoint engagers protect HLA-deficient iPSCs and derivatives from innate immune cell cytotoxicity. Cell Stem Cell. 2023;30(11):1538-1548.e4.

18. Philip B, Kokalaki E, Mekkaoui L, et al. A highly compact epitope-based marker/suicide gene for easier and safer T-cell therapy. Blood. 2014;124(8):1277–1287.

19. Cheng MI, Li JH, Riggan L, et al. The X-linked epigenetic regulator UTX controls NK cell-intrinsic sex differences. Nat. Immunol. 2023;24(5):780–791.

20. Gumá M, Angulo A, Vilches C, et al. Imprint of human cytomegalovirus infection on the NK cell receptor repertoire. Blood. 2004;104(12):3664–3671.

21. Garff-Tavernier ML, Béziat V, Decocq J, et al. Human NK cells display major phenotypic and functional changes over the life span. Aging Cell. 2010;9(4):527–535.

22. Zuhair M, Smit GSA, Wallis G, et al. Estimation of the worldwide seroprevalence of cytomegalovirus: A systematic review and meta-analysis. Rev. Med. Virol. 2019;29(3):e2034.

23. Zhang J, Hu Y, Yang J, et al. Non-viral, specifically targeted CAR-T cells achieve high safety and efficacy in B-NHL. Nature. 2022;609(7926):369–374.

24. Mackensen A, Müller F, Mougiakakos D, et al. Anti-CD19 CAR T cell therapy for refractory systemic lupus erythematosus. Nat Med. 2022;28(10):2124–2132.

