## Supplemental Methods and Figures for "Universal protection of allogeneic cell therapies from natural killer cells via CD300a agonism"

##### *Cell Isolation*

T Cells and NK cells were isolated from fresh Leukopaks sourced from HemaCare and StemCell Technologies. Leukopaks are depleted of CD14+ cells according to manufacturer instructions (Biolegend Cat# 480026). NK cells are isolated from the CD14-depleted fraction by positive selection using anti-CD56 magnetic beads according to manufacturer instructions (StemCell Cat# 17855). T cells were isolated from the CD56-depleted fraction by negative selection according to manufacturer instructions (StemCell Cat# 17951). In some cases, T cells were isolated from buffy coats by density gradient centrifugation according to manufacturer instructions (StemCell Cat# 15061). Primary T and NK cells were cryopreserved at 5e6-20e7 cells/mL using Bambanker (GC Lymphotec BB02).

##### *CRISPR/Cas9 KO and Non-viral HDR T Cell Editing*

Cryopreserved human primary T cells are thawed and rested overnight in Rh10p media containing RPMI (Gibco 61870127), 10% v/v heat-inactivated human AB serum (Gemini Bio 100-512-100, 1x Antibiotic-Antimycotic (Gibco 15240062), and supplemented with 40 ng/mL IL-2 (Peprotech 200-02), 10 ng/mL IL-7 (Peprotech 200-07), and 10 ng/mL IL-15 (Peprotech 200-15) to make Rh10pc. Rested T cells are activated at 1:1 ratio with CD3/CD28 dynabeads (ThermoFisher 11131D) for 2 days. Activated T cells are resuspended to 5-10 million per 100 µL in buffer P3 (Lonza) containing 5 µg HDR template and 2.5 µM of each relevant Cas9 RNP. Cells are electroporated (EP) using Lonza Nucleofector 4D with program EH-115, rested by addition of 900 µL of pre-warmed R10 (RPMI + 10% v/v FBS) media for 30 minutes, and then transferred into one 6-well Grex (Wilson Wolf 80240M) containing 20 mL of warm Rh10pc for overnight rest. Media is replenished the following day and every 2-3 days and split to maintain a cell concentration of ~0.5-2 million/mL hereafter. B2M Knockout (B2M KO) T cells for mRNA electroporation experiments are generated in the same manner as above but without HDR template.

HDR-expressing T cells are enriched 3-4 days post-EP using an appropriate PE-labeled antibody and anti-PE magnetic beads following manufacturer instructions (StemCell Cat# 17684). HLA-A2 specific TCRαβ, HLA-E, NKG2A/CD300a TASR, and SIRPα/TIM3 engager transgenes were enriched using PE-labeled anti-Myc, (R&D IC3696P), anti-HLA-E (Biolegend Cat 342604), anti 218 Linker (CST 62405S), and anti G4S Linker (CST 38907S) antibody, respectively. RQR8-containing HDR templates, such as CD47 and GFP, were enriched using PE-labeled clone Qbend10 antibody (R&D FAB7227P). Magnetically enriched cells are cultured overnight in Rh10pc and then restimulated using Immunocult CD2/CD3/CD28 activator (StemCell 10990) for 7-10 days of further expansion prior to cryopreservation in Bambanker (GC Lymphotec BB02).

Duplex integration of Cloaking into B2M locus and CAR into the TRAC locus was performed as above with the following modifications. 3.75 µg of each HDR template, or 7.5 µg total, is used for EP. Cells are transferred post-EP into Rh10pc media supplemented with 10 µM XL413 (Selleckchem S7547), 0.05 µM Trichostatin A (Selleckchem S1045), and 0.5 µM AZD7648 (Selleckchem S8843); Media is exchanged 1-day post-EP to dilute the small molecule supplements at least 6-fold. Overnight rested, magnetically enriched T cells as above are restimulated using mitomycin-C treated Raji cells at 1:1 ratio of T Cell:Raji Cell. Raji cells were incubated for 1 hour, 37c, at 3 million/mL in R10p (RPMI, 10% v/v FBS, 1x Antibiotic-Antimycotic) supplemented with 50 ug/mL Mitomycin-C (StemCell Cat#100-1048), and then washed four times with R10p.

Alloreactive T cell clones (allo-T) are generated by replacement of the endogenous TCRαβ with a known alloreactive TCRαβ into the TRAC locus and ablation of endogenous TRBC by non-viral HDR. TCRαβ clone AHIII binds to peptide ALWGFFPVL derived from EMC7 protein presented on HLA-A2. TCRαβ clone MJ2-DP19 binds peptide YIGEVLVSV derived from MYO1G protein presented on HLA-A2, otherwise known as minor histocompatibility antigen 2 (HA-2) (patent

WO2022221478A1). A 2xMyc tag was inserted between the signal peptide and variable region of TCRα to enable identification and sorting of transfected T cells. The HDR templates sequences for both TCRs are located in supplementary table 3. T cell clones were generated from HLA-A2 negative donors, with serotyping assessed by flow cytometry of PBMCs stained with PE-labeled anti-HLA-A2 antibody (Biolegend 343306).

*mRNA Construction, Synthesis, and Electroporation*

mRNA-encoding DNA templates are constructed containing a T7 promoter and Human alpha-1-globin 5' Untranslated Region. All TASR and synthetic engager mRNAs are bicistronic with the GFP variant mNeonGreen in the second position connected via T2A linker. mRNA encoding natural ligands are encoded as a single gene. Negative control GFP mRNA was purchased directly (Trilink L-7601-100), while all other mRNA was in-vitro transcribed according to manufacturer instructions and purified with lithium-chloride precipitation (NEB E2060S). The DNA template sequences of all mRNAs are listed in Supplementary Table 4.

For EP, B2M KO T cells are thawed and stimulated with Immunocult CD2/CD3/CD28 activator for 3-4 days in Rh10pc. 1-2 million activated B2M KO T cells are EP'd in 20 µL P3 buffer with code CM-137 using Lonza Nucleofector 4D with 1.5 ug of mRNA, recovered for 15 minutes at 37c after adding 80 µL pre-warmed R10 media, and then plated into 500 µL Rh10pc for overnight rest. For mRNA templates that do not encode GFP, 200 ng of GFP mRNA is added alongside the main mRNA. 1-day post-EP, T cells are assessed for mRNA expression by flow cytometry and for co-culture.

*Allo-T, NK Cell Challenge Assay*

Cryopreserved allo-T cells are thawed, stimulated with Immunocult CD2/CD3/CD28 activator (StemCell 10990), and cultured for 6-7 days in Rh10pc. Cryopreserved NK cells are thawed and cultured in Rh10p + 10 ng/mL IL-15 for 3-5 days prior to start of co-culture. HDR edited T cells are thawed and cultured for 3-4 days in Rh10pc. For co-cultures, 30,000 HDR edited T cells are plated with NK and/or allo-T cells at various E:T ratios in a 96 well round-bottom plate for 20 hours in R10p + 10 ng/mL IL-15.

Co-cultures were subjected to Fluorescent Barcoding Flow Cytometry using the following protocol. Amine-reactive dyes AF647 (Thermofisher A20006), IF700 (AATBioquest 71514), and IR800CW (Licor 929-70021) are resuspended in DMSO and mixed in 8 combinations, described in Supplemental Fig. 1, to make 100X working stocks of 0.2 mg/ml, 1 mg/ml, and 1.5 mg/ml, respectively, and stored in -80C. The co-culture plate was resuspended to 50 µL PBS by centrifugation at 500g for 2 minutes to decant supernatant. Fluorescent barcode working stocks were thawed, diluted 1:50 with PBS, and 50 µL of the appropriate barcode combination was immediately transferred to the corresponding wells of the co-culture plate using a multichannel pipette and incubated in the dark for 15 minutes room temperature. 100 µL of R10p is added to each well and incubated for 10 minutes to quench the reaction, followed by two washes in FACS buffer by centrifugation at 1200g for 2 minutes. Wells are mixed column-wise and transferred into a new 96-well round-bottom plate, spun 1200g 2 minutes, and then resuspended in the following antibody staining mix in FACS buffer: BV421 CD56, BV510 CD3, BV605 HLA-I, 7-AAD, PE Qbend10. The plate is washed twice in FACS buffer by centrifugation at 1200g for 2 minutes, and then acquired at equal volumes on flow cytometry.

*PBMC Challenge Assay*

AAVS1 HDR edited T cells are thawed and stimulated with Immunocult CD2/CD3/CD28 activator for 6-7 days in Rh10pc. Cloaking expression levels are assessed on the day of co-culture. Cryopreserved PBMCs are sourced from Hemacare and StemCell Technologies. PBMCs are thawed, assessed for phenotype by flow cytometry, and rested overnight in Rh10p + 40 ng/mL IL-2. For co-culture, 30,000 HDR edited T cells are plated with PBMCs at various E:T ratios in a 96 well flat-bottom plate for 3 days in R10p + 40 ng/mL IL-2. Cells are transferred to a 96 well round-bottom plate, and T cell survival is determined by fluorescent barcoding flow cytometry using the protocol outlined in the Allo-T cell challenge assay but using the following antibody staining mix: PE Qbend10, BV605 HLA-I, BV650 CD56, BV785 CD3, 7-AAD.

*Allo-T + NK Cell Competition Assay*

Allo-T cells and NK cells are thawed and cultured as per the Allo-T and NK challenge assay protocols above. B2M HDR edited T cells are thawed and cultured for 3-4 days in Rh10pc, and then pooled together at 1:1 ratio. Pooled HDR edited T cells are seeded at 200,000 per well and co-cultured with Allo-T and NK cells at various E:T ratios in a 96 well flat-bottom plate for 20 hours in R10p + 10 ng/mL IL-15. 10 µL of counting beads (Biolegend 424902) are added to each well and transferred to new 96 well round-bottom plate. The plate is spun 500g 3 min and resuspended into 50 µL of antibody staining mix, consisting of the following reagents in FACS buffer: BV421 Qbend10, BV510 CD3, BV605 HLA-I, BV785 CD56, PE 218 Linker, 7-AAD, PE-Cy7 HLA-E, AF647 G4S Linker, and APC-Cy7 CD47. After staining, cells washed twice with FACS buffer and acquired on Flow cytometry at equal volumes per well.

*B Cell Depletion Assay*

Cryopreserved, HDR edited CAR-T cells are thawed and cultured for 3-4 days in Rh10pc. Cryopreserved PBMCs are thawed and cultured in Rh10p + 40 ng/mL IL-2 for 3 days. PBMCs are seeded at 250,000 per well and co-cultured with CAR-T cells at the indicated CAR-T:PBMC ratio in a 96 well flat-bottom plate for 3 days in R10p + 40 ng/mL IL-2.

B cell depletion is determined using the following antibody-based fluorescent barcoding flow cytometry protocol. Cells are transferred to a 96 well round-bottom plate spun by centrifugation at 500g for 3 minutes, and then resuspended into 50 µL of antibody and antibody-based barcode staining mix. The following antibody staining mix in 30µL of FACS buffer: BV421 CD56, BV510 CD3, PE CD19, AF647 G4S Linker, 7-AAD are first added, and then 20µL of the antibody-based barcode mix are added. Barcode staining mix consists of a combination of anti-CD45 antibodies on the BV711, BV785, PE-Cy7, and APC-Cy7 Channels in FACS buffer that are unique to each row of the plate, as described in Supplemental Fig. 12. The plate is washed twice in FACS buffer by centrifugation at 500g for 3 minutes. Wells are mixed column-wise and transferred into a 96-well deepwell plate and then acquired at equal volumes on flow cytometry.

*Cloaking Ligand Expression by Flow Cytometry*

Expression of cloaking ligands on T cells was assessed by flow cytometry using antibody-based and ligand-based staining. Expression of TASRs were first assessed by staining with 5 ug/mL of FC-tagged recombinant protein ligands followed by PE anti-FC secondary antibody. In addition, expression of NKG2A TASR, TIM3 engager, and SIRPa engager determined by staining with 5 ug/mL of biotinylated recombinant proteins NKG2A, TIM3, and SIRPα, respectively, followed by PE-streptavidin secondary staining. CD300a and NKG2A TASR expression also determined using PE labeled anti-218 Linker antibody. Anti-CD19 CAR, TIM3 engager, and SIRPa engager expression also determined using PE and AF647 labeled anti-G4S linker antibody. HLA-E expression is determined using BV421, PE, or PE-Cy7 labeled anti-HLA-E antibody. HLA-G expression is determined using PE labeled anti-HLA-G antibody. CD47 expression determined using BV421, PE, or APC-Cy7 labeled anti-CD47 antibody. A list of all reagents can be found in Supplementary Table 1.

*Raji Cytotoxicity*

Cryopreserved, HDR edited CAR-T cells are thawed and cultured for 5-6 days in Rh10pc. Cryopreserved Luciferase-expressing Raji cells expressing are thawed and cultured for 5-6 days in R10p. 20,000 Raji cells are seeded per well and co-cultured with CAR-T cells at various E:T ratios in a 96 well opaque flat-bottom plate for 20 hours in R10p. Raji cell cytotoxicity is determined by luciferase expression using a luminescence plate reader according to manufacturer instructions (Promega E6120).

#### Supplemental Tables 1-4

**Supplemental Table 1: Antibody panels and protein staining reagents for flow cytometry readouts**

**Supplemental Table 2: List of gRNA sequences**

**Supplemental Table 3: HDR template sequences used to generate engineered T cells by non-viral HDR**

**Supplemental Table 4: mRNA-encoding DNA template sequences**

### Supplemental Figures 1-13

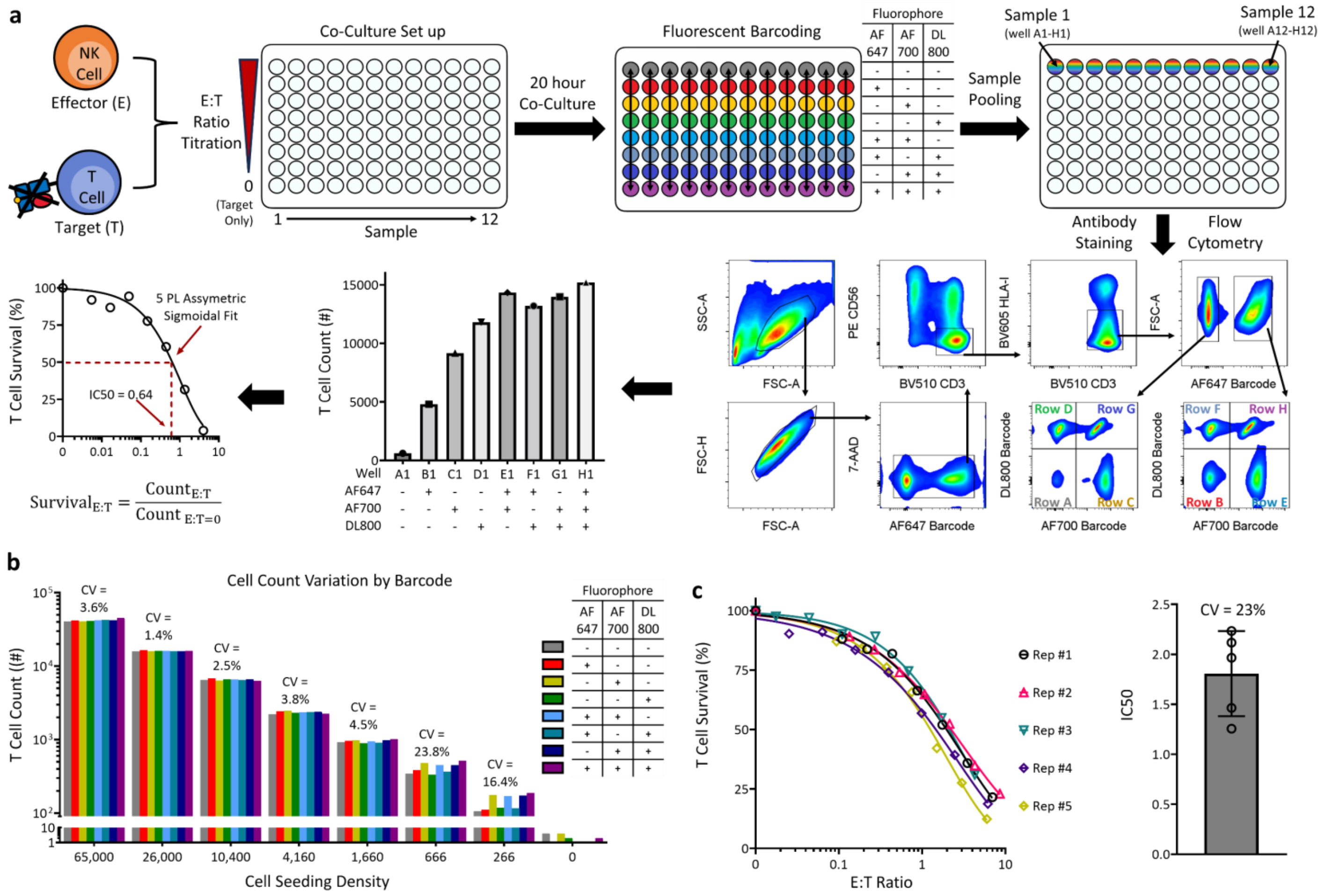


**Supplemental Figure 1: Experimental overview of T cell survival assay against allogeneic cell challenge by fluorescent barcoding flow cytometry.**

**a**. Experimental overview. T cell targets are plated with allogeneic effectors, such as NK cells, using the plate map indicated, where each column represents a unique pair of target and effectors at varying E:T ratios with the target cell count fixed. After co-culture, every row is stained with a unique combination of three amine-reactive fluorophore dyes as indicated. Each column of 8 wells is then pooled into a single well, stained with antibody, and then run on flow cytometry. Target T cells are first gated, followed by barcode deconvolution. In cases where T cell targets do not have B2M KO, the HLA-I- gate is not used. The percent survival of target cells from a given well, corresponding to one E:T ratio, is calculated by normalizing over the target only well(s). The resulting curve of T cell survival versus E:T ratio is fit to the 5-parameter logistic function with maximum and minimum constraints set to 100% and 0% respectively. IC50 value represents the E:T ratio corresponding to 50% target T cell survival. Each column on the plate map represents one technical replicate curve. **b**. T cells were seeded at the indicated counts to all wells of a given column on a 96-well plate and then subjected to cell counting by fluorescent barcoding flow cytometry. Every group represents one column from seeded at the indicated number of T cells per well. Subgroups represent the cell count corresponding to the given barcode for the indicated group. CV indicates coefficient of variation for the indicated group, since cell counts should be the same for all barcodes within a group. **c**, Survival curves of B2M KO T cells EP'd with GFP mRNA challenged with one NK cell donor. The same experiment was repeated 5 times on different days. (right panel) CV of IC50 value. N = 1 technical replicate curves per experiment

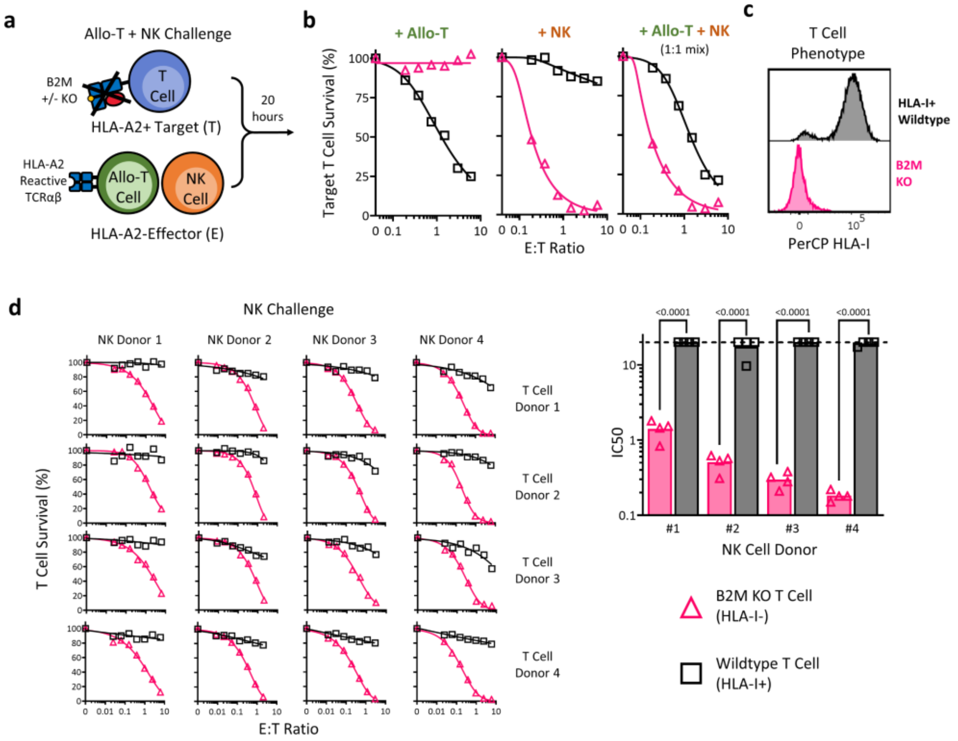


**Supplemental Figure 2: B2M KO rescues allogeneic T cells from T cell alloreactivity but introduces NK cell alloreactivity**

**a**, Experimental overview. Survival of Wildtype (WT) and B2M KO T cells from an HLA-A2+ donor challenged with NK and allo-T cell effectors from HLA-A2- donors. Allo-T cells contain the AHIII T cell receptor reactive to EMC7 peptide, ALWGFFPVL, presented by HLA-A2. B2M KO T cells express RQR8 epitope tag from the B2M locus, WT T cells express GFP-2A-RQR8 from the AAVS1 locus under control of EF1a promoter. **b**, Survival curves of WT and B2M KO target T cells challenged with the indicated effectors. Target T cells are specifically gated by CD3+CD56-RQR8+. N = 1 technical replicate curves per condition. **c**, HLA-I expression of WT and B2M KO T cells by flow cytometry, gated live single cell lymphocytes. (d) B2M KO specifically renders T cells susceptible to NK cell rejection. **d**, NK challenge assay with multiple NK donors and multiple target T cell effectors. Bar graph represents summary of IC50 value, each data point represents WT or B2M KO T cell from one T cell donor against the indicated NK cell donor. Mann Whitney U-Test.

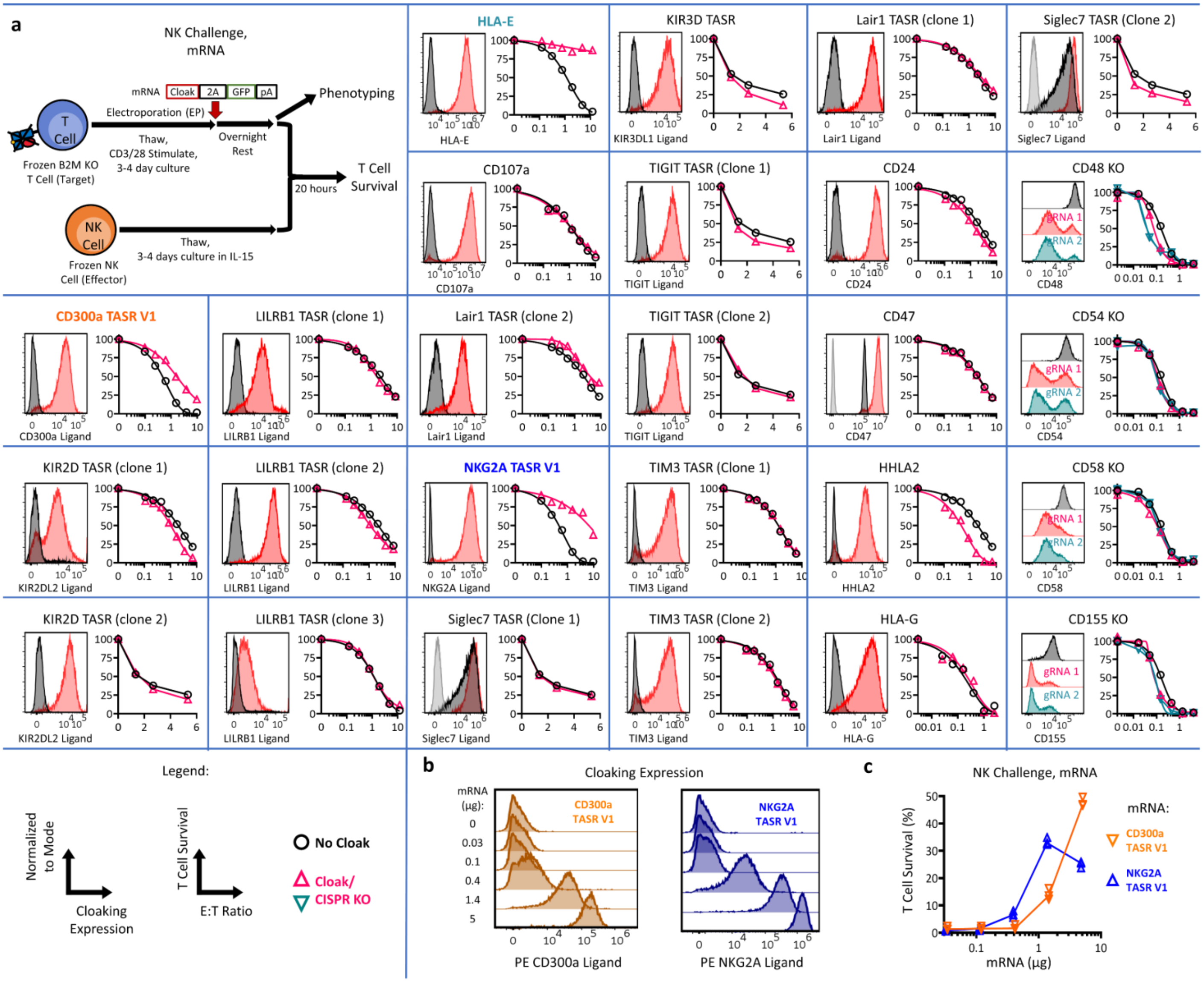

**Supplemental Figure 3: Functional screening and validation of NK cloaking strategies by mRNA electroporation and CRISPR KO of B2M KO T cells, related to Figure 1a**

**a,** (top left) Process diagram for expression of cloaking ligands by mRNA EP and functional validation. Each inset represents a single screen member. CRISPR KO are in addition to B2M KO. Negative control, in black, is GFP mRNA for mRNA EP conditions or B2M KO only for the CRISPR KO screen. (Left inset plot) expression profile by flow cytometry, gated single cell lymphocytes. Red and Teal indicate the given ligand, black indicates negative control. Grey on Siglec7 and CD47 indicates unstained. TASR expression determined by staining with recombinant cognate ligand. (Right inset plot) NK challenge assay of the given screen member. **b-c**, validation of CD300a TASR V1 and NKG2A TASR V1 by mRNA titration. (b) Expression of NKG2A and CD300a TASR by flow cytometry on B2M KO T cells 1-day post electroporation (EP) with the given quantity of respective mRNA. (c) Survival of B2M KO T cells electroporated CD300a or NKG2A TASR at the indicated mRNA dose as in (b) challenged with NK cells at a constant E:T ratio of 4.6. N = 3 technical replicates per condition.

a

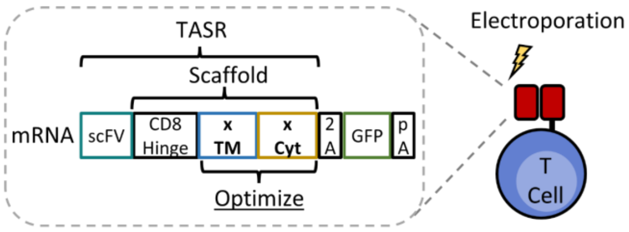


b

| TM+Cyt Name | TM Sequence | Cyt Sequence |
|---|---|---|
| CD8 TM (V1) | IYIWAPLAGTCGVLLLSLVITLYCN | |
| mB7-1 TM+Cyt | PPEDPPDSKNTLVLFGAGFGAVITVVVIVVII | KCFCKHRSCFRRNEASRETN NSLTFGPEEALAEQTVFL |
| KIR2DL1 TM | PRHLHILIGTSVVIILFILLFFLLHRWCSNKKNAA VMDQ | |
| HLA-C06 TM+Cyt | VGIVAGLAVLAVLAVLGAVMAVVMC | RRKSSGGKGGSCSQAASSNS AQGSDESLIACKA |
| proCAR-3 TM | PFWLLFILVAILGGLFGAIVAFLLALWVR | |
| HLA-b57 TM+Cyt | VGIVAGLAVLAVVVIGAVVAAVMC | RRKSSGGKGGSYSQAACSDS AQGSDVSLTA |
| proCAR-4 TM | PFWLLVALLALLAVIAALLAAIFALLWVR | |
| ITGA TM | VPLWVILLSAFAGLLLLMLLILALW | |

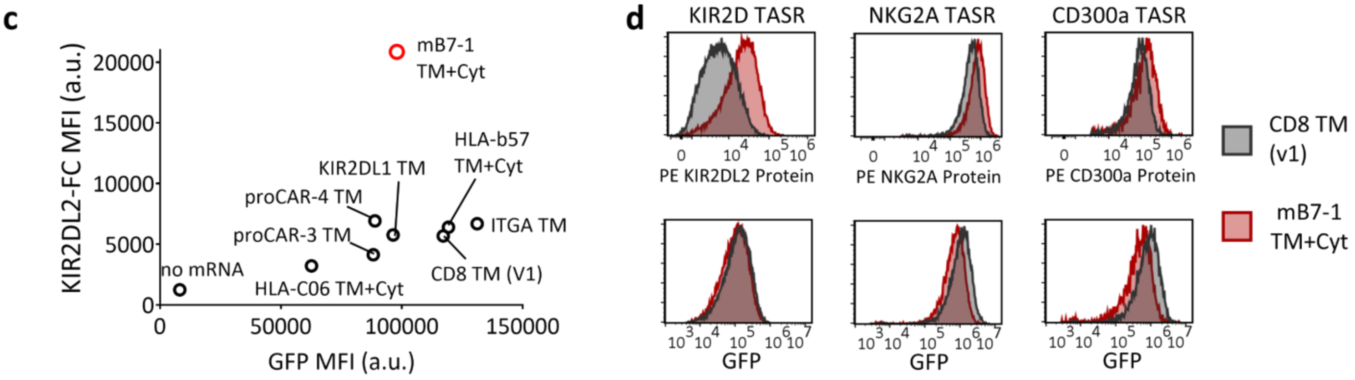


**Supplemental Figure 4: Mouse B7-1 domains enhance TASR expression**

**a**, Experimental overview. TASRs were constructed with various transmembrane and cytoplasmic domains, in-vitro transcribed into mRNA, and then transiently transfected into primary T cells to test for expression. All constructs were bicistronic with GFP to enable normalized comparison between difference constructs. **b**, various TM+Cyt domains tested, CD8 TM is from the V1 scaffold. **c**, GFP versus TASR median fluorescence intensity (MFI) of a KIR2D specific TASR with the various TM+Cyt domains from (b) as assessed by flow cytometry, gated single cell lymphocytes. mB7-1 provides higher TASR expression normalized to GFP translation. **d**, Expression of KIR2D, NKG2A, and CD300a TASRs with either CD8TM or mCD80 TM+Cyt after mRNA EP as measured by flow cytometry, gated single cell lymphocytes.

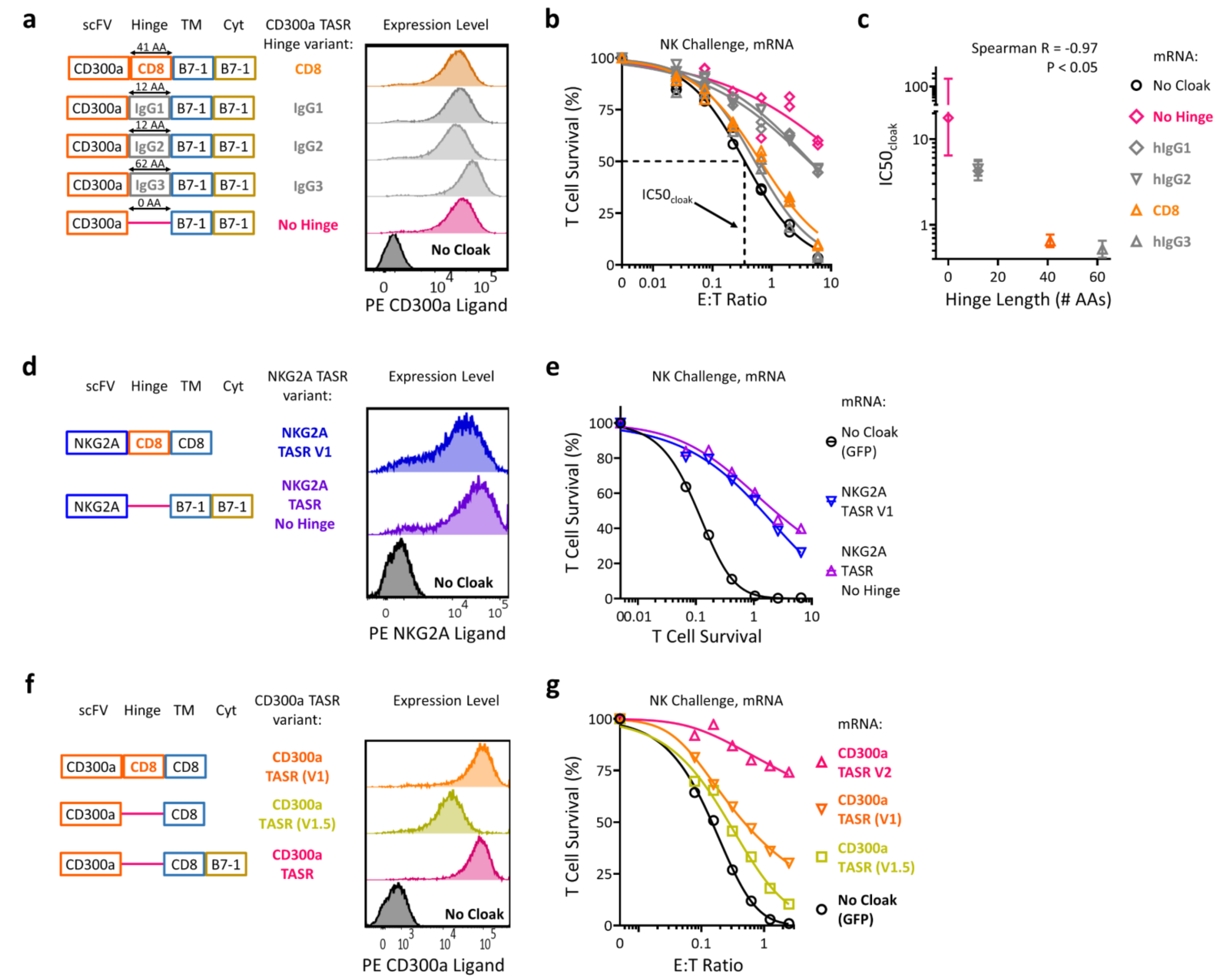


**Supplemental Figure 5: Inverse correlation of hinge length and functional potency of CD300a TASR but not NKG2A TASR, and final optimization of CD300a TASR (V2)**

(a-e), effect of hinge length on function of CD300a TASR and NKG2A TASR. **a**, structure of CD300a TASR variants with different hinge domains with the indicated amino acid lengths and their expression level after mRNA EP into B2M KO T cells. **b**, NK challenge assay of B2M KO T cells expressing the indicated CD300a TASR variants. N = 2 technical replicate curves per condition. **c**, correlation of NK protection from (b) as defined by IC50 value with hinge length. Error bar indicates 95% confidence interval of IC50 value. **d**, structure of NKG2A TASR V1 and a no hinge variant, along with their expression level by mRNA EP into B2M KO T cells. **e**, NK challenge assay of B2M KO T cells expressing the indicated NKG2A TASR variant, N = 1 technical replicate curve per condition. (f-g), Final optimization of CD300a TASR. **f**, CD300a TASR V1 and two optimized variants are tested for expression in via mRNA EP of B2M KO T cells. **g**, NK challenge assay of B2M KO T cells expressing the indicated CD300a TASR variant, N = 1 technical replicate curve per condition.

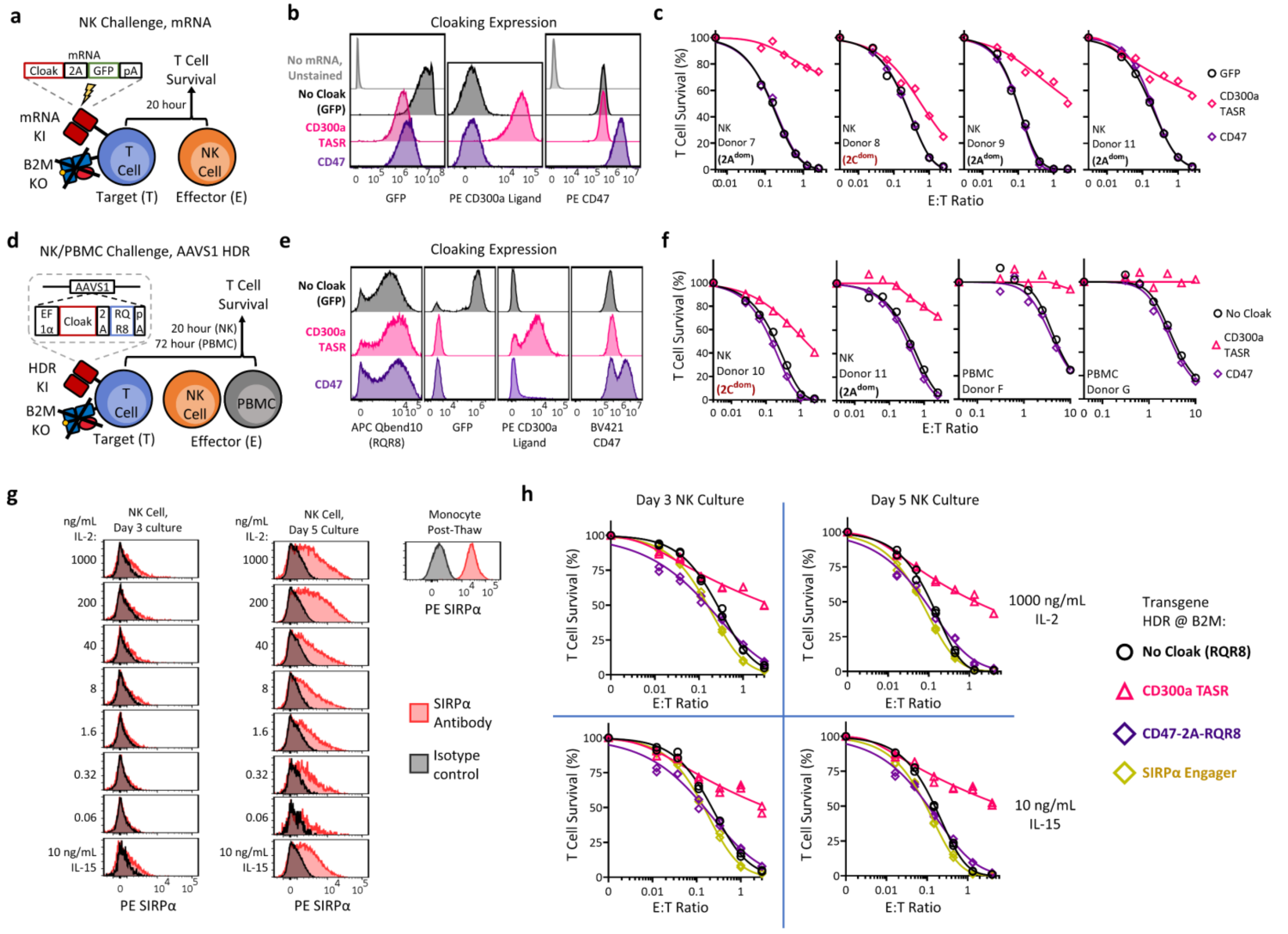


**Supplemental Figure 6: CD300a TASR outperforms CD47 in B2M KO T cells in additional model systems and with IL-2 activated NK cells**

(a-c) Comparison of CD300a TASR and CD47 via mRNA electroporation. **a**, experimental overview. **b**, Flow cytometry phenotype of T cells electroporated with the indicated mRNA. **c**, NK challenge assay of T cells in (b) against four NK cell donors with either $2A^{dom}$ or $2C^{dom}$ NK phenotype. N = 1 technical replicate curves per condition. (d-f) Comparison of CD300a TASR and CD47 expressed from the AAVS1 loci of B2M KO T cells in both NK and PBMC challenge assays. **d**, experimental overview. **e**, Flow cytometry phenotype of T cells containing the indicated cloaking transgene or GFP, gated single cell lymphocytes. **f**, NK and PBMC challenge assay of T cells from with either NK or PBMC donors as indicated. N = 1 technical replicate curves per condition. **g**, Effect of cytokine concentration and culture time on the expression of SIRPα on cultured human NK cells. Cryopreserved NK cells were thawed and cultured at the indicated concentrations of IL-2 or 10 ng/mL IL-15, used for the standard NK challenge assay, for 3 and 5 days. Monocytes served as positive staining control. NK cells are gated CD3-CD56+, Monocytes are gated CD14+$FSC^{hi}SSC^{hi}$. **h**, NK challenge assay with B2M KO T cells expressing the indicated cloaking transgene at B2M loci as in Fig. 1c. NK cells were cultured for the indicated duration with the indicated cytokine, during both initial culture as in (g) and during the 20-hour co-culture. N = 2 technical replicate curves per condition.

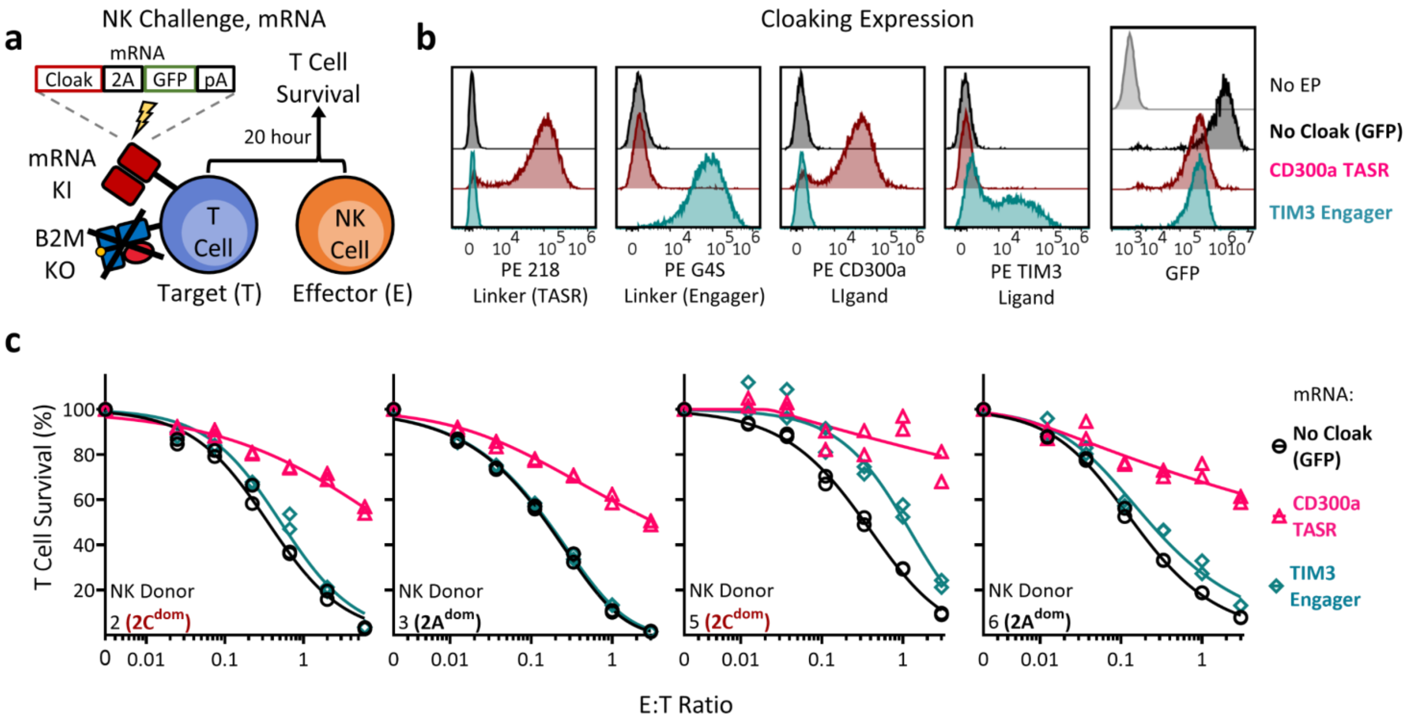


**Supplemental Figure 7: CD300a TASR outperforms TIM3 Engager**

**a**, experimental overview for comparison of CD300a TASR with TIM3 engager via mRNA electroporation. Both ligands were bicstronic with GFP. **b**, Cloaking ligand expression assessed using both ligand-based staining and antibody-based staining by flow cytometry with the indicated mRNA, gated single cell lymphocytes. **c**, NK challenge assay of T cells from (b) challenged with the four indicated NK cell donors. N = 2 technical replicate curves per condition.

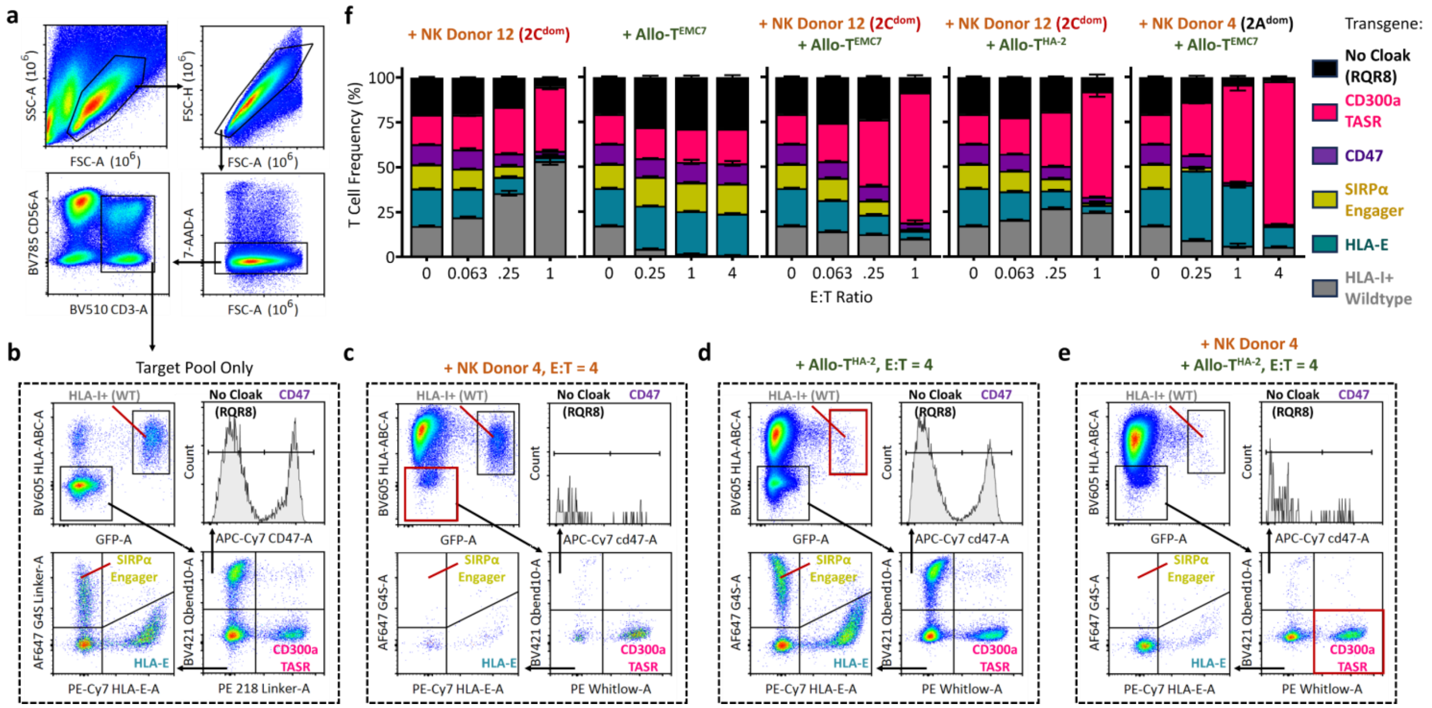


**Supplemental Figure 8: Flow cytometry gating scheme and additional conditions for Allo-T + NK competition assay, related to Fig. 1e.**

(a-e) representative gating strategy and readout for competition assay. All samples were resuspended and acquired at equal volumes. **a**, pre-gating on CD3+ single-cell lymphocytes. **b**, gating for target pool only with no effector cells. Labels designate the pool member. The WT T cell member express GFP and RQR8 from the AAVS1 locus. HLA-I+GFP- cells represent residual cells from incomplete B2M KO T cells. **c**, gating with NK effector challenge only, red gate highlights depleted population relative to (b). **d**, gating with allo-T cell effector challenge only, red gate and label highlights depleted population relative to (b). **e**, gating with allo-T and NK cell challenge, red gate highlights CD300a TASR survival relative to other members and (b). **f**, additional NK and Allo-T challenge conditions as in Fig. 1. Allo-T$^{EMC7}$ contains the AHIII T cell receptor reactive to EMC7 peptide, ALWGFFPVL, presented by HLA-A2. NK and allo-T cells effectors are mixed 1:1. N = 3 technical replicates per condition.

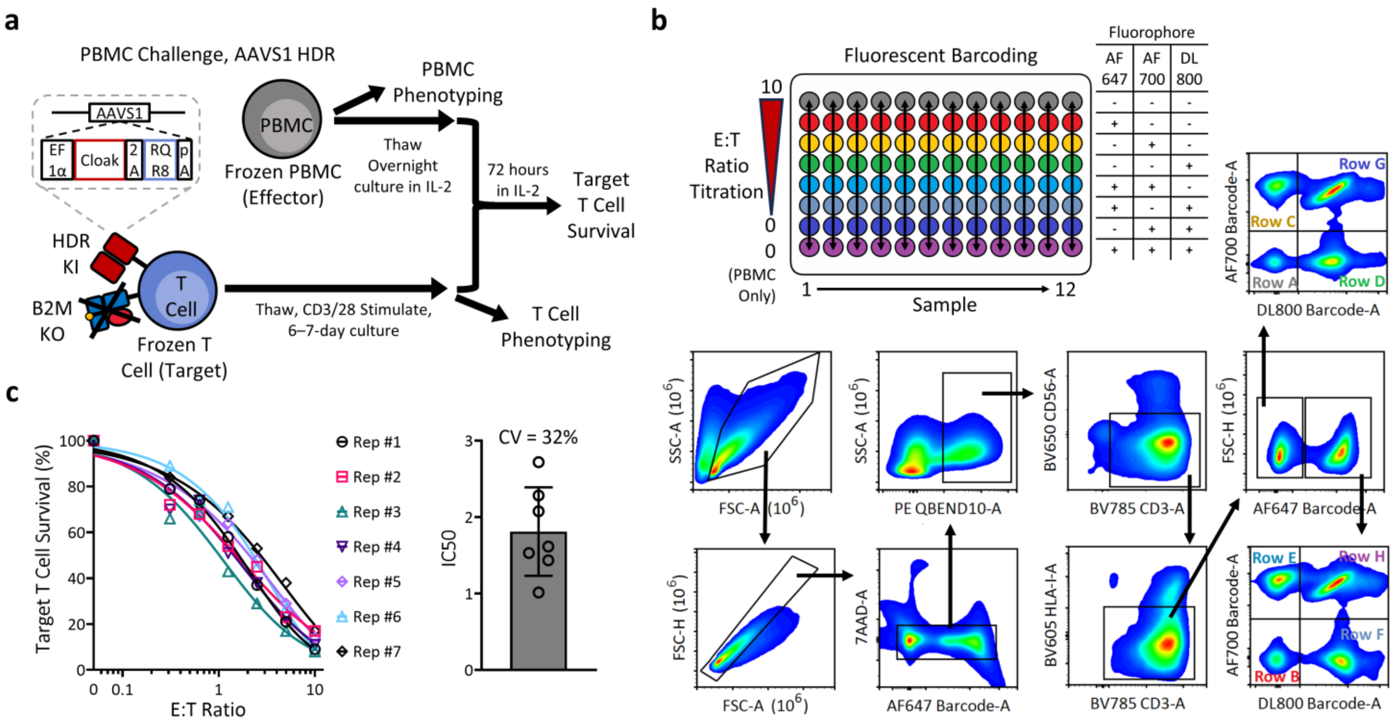


**Supplemental Figure 9: Experimental overview of PBMC challenge assay and validation, related to Fig. 2a-f**

**a**, Experimental overview of PBMC challenge assay. PBMC phenotyping was performed post-thaw, and cloaked T cell phenotyping performed at time of co-culture. **b**, co-culture plate map and gating scheme for fluorescent barcoding flow cytometry readout in the PBMC challenge assay. **c**, Inter-assay variation of PBMC challenge assay of one PBMC donor against one B2M KO T cell source. Experiments performed on separate days, N = 1 technical curve per experiment. (right panel) CV of IC50 values.

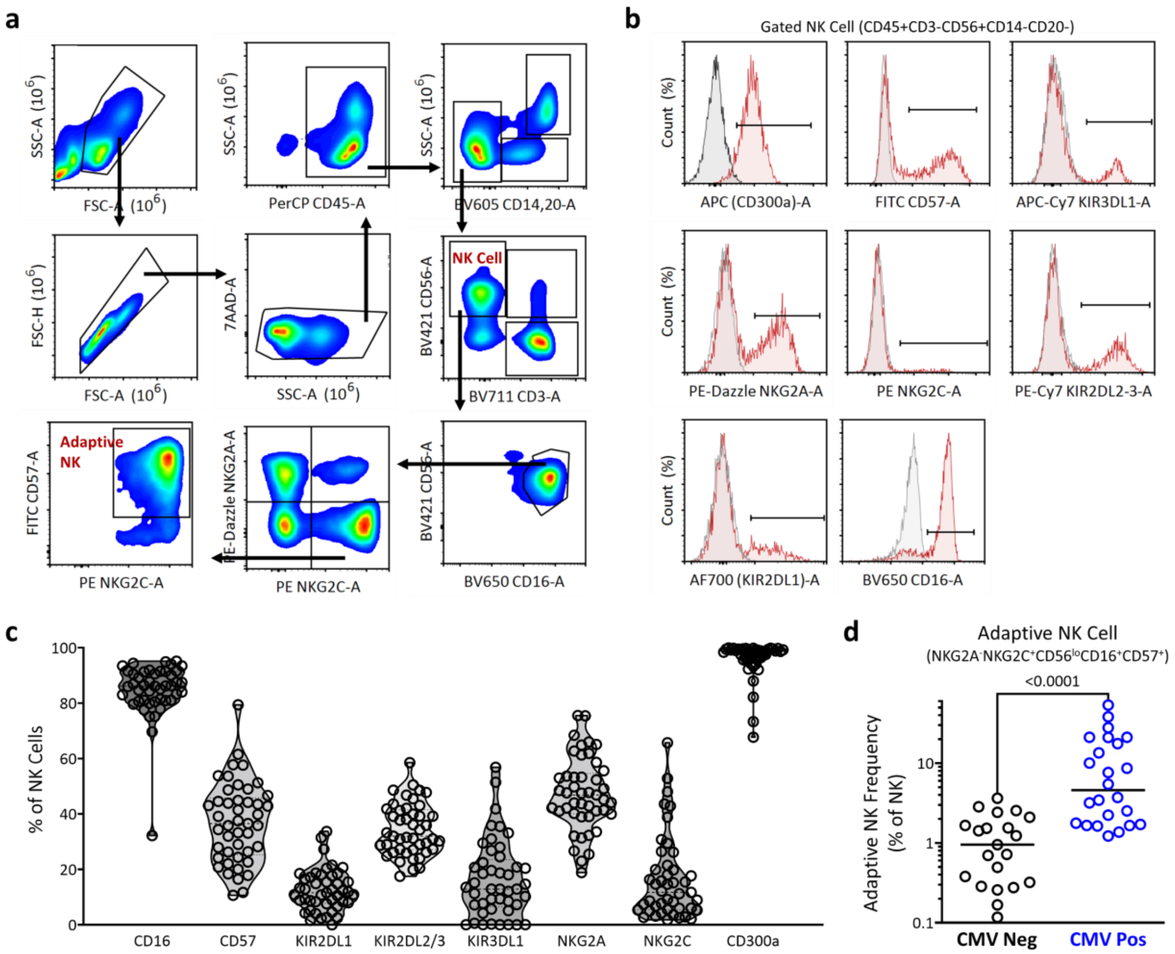


**Supplemental Figure 10: NK cell phenotyping of PBMC challenge assay, related to Fig. 2a-f**

**a**, gating scheme for NK cells and adaptive NK cell phenotype from PBMC. **b**, representative expression of NK cell phenotyping markers from one PBMC donor. Red indicates full panel staining, grey indicates CD300a fluorescence-minus-one negative control staining. **c**, Percentage of NK cells expressing the indicated marker with gating from (b). N = 45 PBMC donors. **d**, Adaptive NK cell frequency gated as in (a) between CMV seropositive (N = 21 donors) and seronegative donors (N = 24 donors), Mann Whitney U Test.

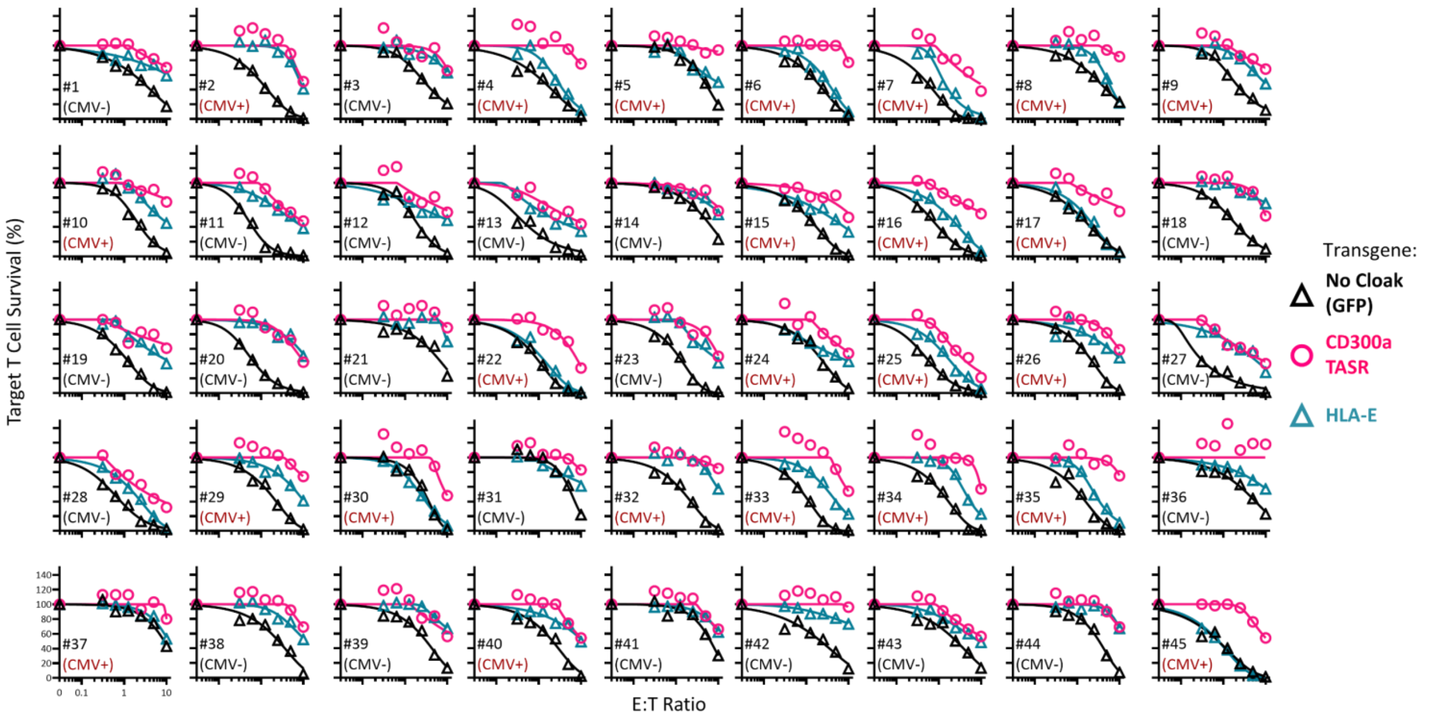


**Supplemental Figure 11: Survival curves of 45 donor PBMC challenge assay, related to Fig. 2d**

Number indicates PBMC donor. Brackets indicate the CMV serostatus of the donor. N = 1 technical replicate curves per condition.

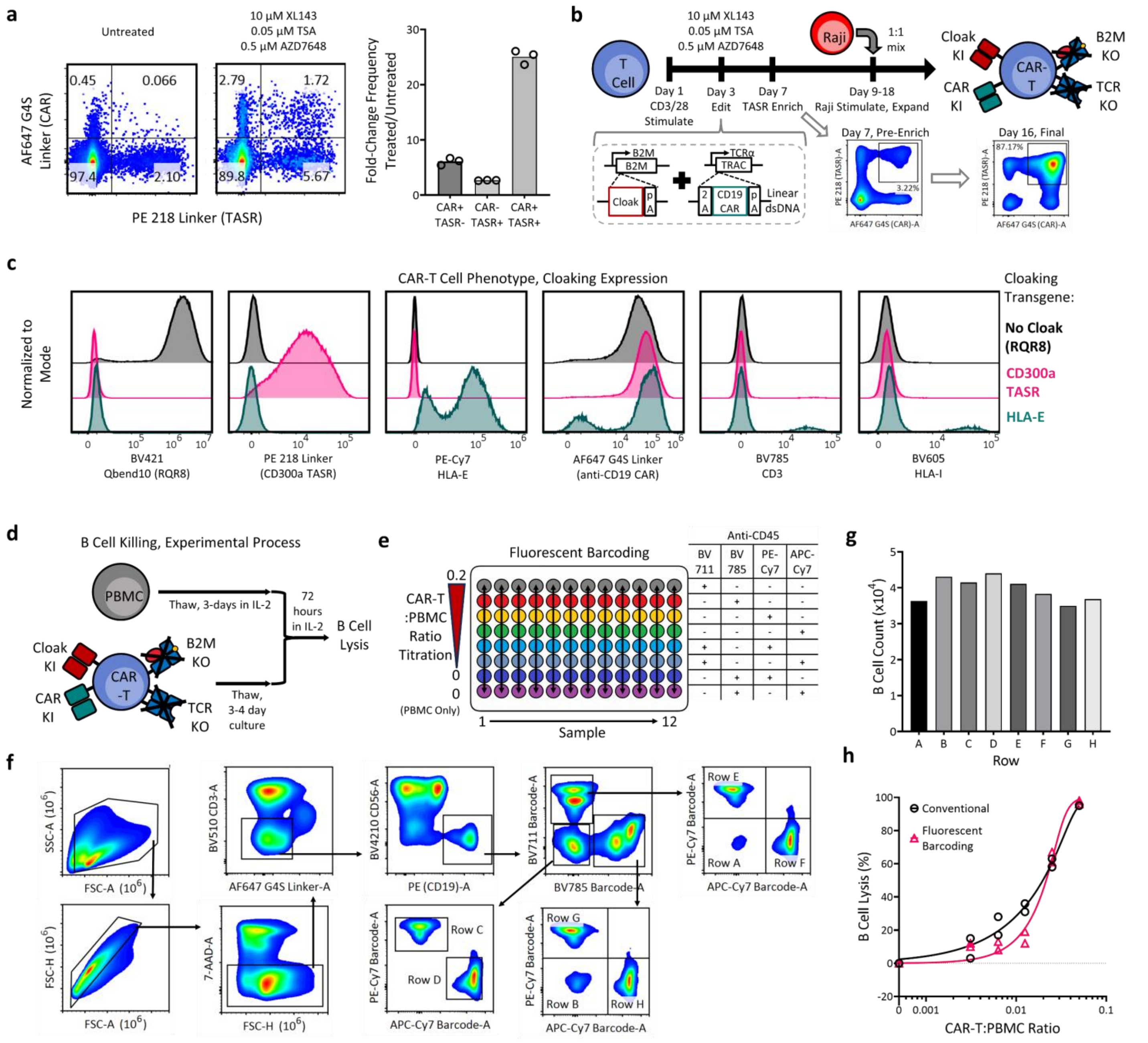

**Supplemental Figure 12. Generation of TASR-expressing CAR-T cells by multiplexed non-viral HDR into primary T cells and use in B cell killing assay, related to Fig. 2g**

**a**, Knock-in efficiency of T cells 4-days post-editing with CD300a TASR at B2M loci and anti-CD19 CAR at TRAC loci. Post-electroporation, T cells were plated into either standard media or media containing the indicated small molecules, dubbed TMX, for 24 hours prior to exchange back into standard media. (right panel) fold enhancement in editing efficiency plated into TMX relative to standard media for CAR+TASR-, CAR-TASR+, and CAR+TASR+ gates in (left panel). N = 3 editing experiments. **b**, Process overview for generation, purification, and expansion of cloaked CAR-T cells. Stimulated T cells are edited in a single step with B2M and TRAC RNPs and linear dsDNA HDR templates encoding cloaking ligand and anti-CD19 CAR. The cloaking HDR template integrates at the start codon of B2M gene while the CAR integrates into TRAC in a bicistronic format via a 2A cleavable peptide. Both constructs code for BGH polyA tail. Cloak ligand expressing cells are enriched using an appropriate antibody by magnetic enrichment, followed by selective expansion of CAR expressing cells by addition of mitomycin-C treated Raji feeder cells. **c**, phenotype of three engineered CAR-T cells by flow cytometry expressed the indicated cloaking transgene at B2M locus, gated live single cells. The same anti-CD19 CAR is used for all CAR-T cells. **d**, Experiment overview for B cell killing assay. **e**, Co-culture plate map and fluorescent barcoding scheme. PBMCs are seeded at 250,000 per well, and then edited CAR-T cells are added at the indicated CAR-T:PBMC ratio. Every column represents one unique PBMC : CAR-T cell pair and cytotoxicity curve. At the end of co-culture, rows are barcoded by staining with unique combinations of fluorescent anti-CD45 antibody along with phenotyping antibodies. The plate is washed and then every column is pooled into one well and acquired on flow cytometry. **f**, gating scheme for identification of B cells and barcode demultiplexing. **g**, B cell counts in the absence of CAR-T cells. PBMCs are seeded at equal density in one column, fluorescently barcoded, and then counted on flow cytometry as in (e,f). **h**, Comparison of B cell lysis measurements by conventional and fluorescent barcoding flow cytometry. RQR8-expressing CAR-T cells were co-cultured with PBMCs at the indicated ratios in four identical columns on the plate. Two columns underwent fluorescent barcoding flow cytometry as in (e,f), while two other columns were directly stained with phenotyping antibodies and then acquired on flow cytometry without pooling. N = 2 technical replicates per kill curve.

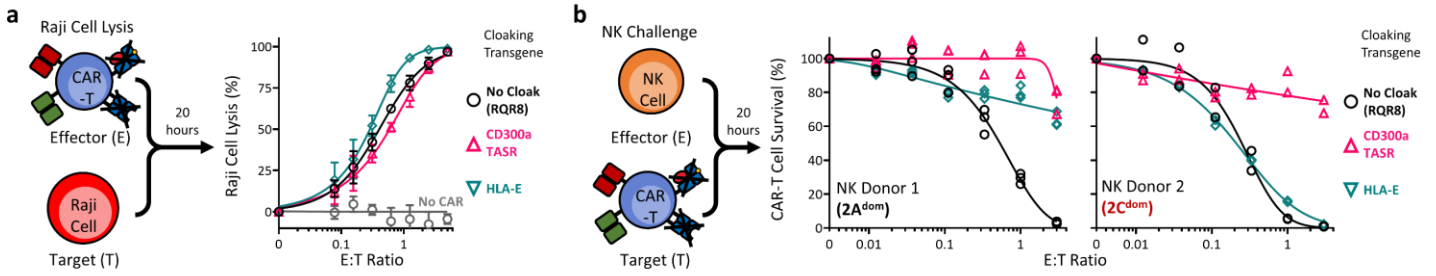


**Supplemental Figure 13. CD300a TASR expressed in a model allogeneic anti-CD19 CAR-T cell exhibit enhanced protection from NK cells with no effect on CAR-mediated killing potency, related to Fig. 2g**

**a**, Cytotoxicity of indicated cloaked anti-CD19 CAR-T cell against CD19-expressing Raji cells. Grey negative control contains CD300a TASR integrated into the B2M loci without anti-CD19 CAR. N = 3 technical replicate curves per condition. **b**, NK challenge of cloaked anti-CD19 CAR-T cells containing the indicated cloaking transgene with a $2A^{dom}$ and $2C^{dom}$ NK donor. N = 3 technical replicates curves per condition.
